# Global diversity and evolution of *Salmonella* Panama, an understudied serovar causing gastrointestinal and invasive disease worldwide: a genomic epidemiology study

**DOI:** 10.1101/2024.02.09.579599

**Authors:** Caisey V. Pulford, Blanca M. Perez-Sepulveda, Danielle J. Ingle, Rebecca J. Bengtsson, Rebecca J. Bennett, Ella V. Rodwell, Maria Pardos de la Gandara, Charlotte E. Chong, P. Malaka De Silva, Magali Ravel, Véronique Guibert, Elisabeth Njamkepo, Neil Hall, Marie A. Chattaway, Benjamin P. Howden, Deborah A Williamson, Jay C. D. Hinton, François-Xavier Weill, Kate S. Baker

**Affiliations:** Clinical Infection, Microbiology, and Immunology, Institute of Infection, Veterinary and Ecological Sciences, University of Liverpool, Liverpool, United Kingdom; Department of Microbiology and Immunology, The University of Melbourne at The Peter Doherty Institute for Infection and Immunity, Melbourne, Australia; Gastrointestinal Bacteria Reference Unit, UK Health Security Agency, London, United Kingdom; Institut Pasteur, Université Paris Cité, Unité des Bactéries pathogènes entériques, Paris, France; Department of Genetics, University of Cambridge, Cambridge, United Kingdom; Earlham Institute, Norwich Research Park, Norwich, UK; Microbiological Diagnostic Unit Public Health Laboratory, Department of Microbiology and Immunology University of Melbourne at The Peter Doherty Institute for Infection and Immunity, Melbourne, VIC, Australia; Centre for Pathogen Genomics, The University of Melbourne, Melbourne, Australia

## Abstract

**Background:** Nontyphoidal *Salmonella* (NTS) is a globally important bacterial pathogen, typically associated with foodborne gastrointestinal infection. Some NTS serovars can also colonise normally sterile sites in humans to cause invasive NTS (iNTS) disease. One understudied *Salmonella enterica* serovar which is responsible for a significant number of cases of iNTS disease is Panama. Despite global dissemination, numerous outbreaks, and a reported association with iNTS disease, *S. enterica* serovar Panama (*S.* Panama) has not been investigated in detail.

**Methods:** Using combined epidemiological and whole genome sequencing data we analysed 836 *S.* Panama genomes derived from historical collections, national surveillance datasets, and publicly available data. The collection represents all inhabited continents and includes isolates collected between 1931 and 2019. Maximum likelihood and Bayesian phylodynamic approaches were used to determine population structure & evolutionary history, and to infer geo-temporal dissemination. A combination of different bioinformatic approaches utilising short-read and long-read data were used to characterise geographic and clade-specific trends in antimicrobial resistance (AMR), and genetic markers for invasiveness.

**Findings:** We identified the presence of multiple geographically linked *S.* Panama clades, and regional trends in antimicrobial resistance profiles. Most isolates were pan-susceptible to antibiotics and belonged to clades circulating in the United States of America, Latin America, and the Caribbean. Multidrug resistant (MDR) isolates in our collection belonged to two phylogenetic clades circulating in Europe and Asia/Oceania, which exhibited the highest invasiveness indices based on the conservation of 196 extra-intestinal predictor genes.

**Interpretation:** This first large-scale phylogenetic analysis of *S.* Panama revealed important information about population structure, AMR, global ecology, and genetic markers of invasiveness of the identified genomic subtypes. Our findings provide an important baseline for understanding *S.* Panama infection in the future. The presence of MDR clades with an elevated invasiveness index should be monitored by ongoing surveillance as such clades may pose an increased public health risk.

**Funding:** The project was supported by both a Global Challenges Research Fund (GCRF) data & resources grant BBS/OS/GC/000009D and the BBSRC Core Capability Grant to the Earlham Institute BB/CCG1720/1. JCDH received a Wellcome Trust Investigator award (grant number 222528/Z/21/Z). CVP was supported by the John Lennon Memorial Scholarship from the University of Liverpool and a Fee Bursary Award from the Institute of Integrative Biology at the University of Liverpool. FXW was awarded the *Institut Pasteur*; *Santé publique France*; the *Fondation Le Roch-Les Mousquetaires*; and the French government’s *Investissement d’Avenir* programme*, Laboratoire d’Excellence* “Integrative Biology of Emerging Infectious Diseases” (grant number ANR-10-LABX-62-IBEID). MDS was supported by a BBSRC grant (KSB: BB/V009184/1). RJBengtsson was supported by an MRC grant (KSB: MR/R020787/1). RJBennett was supported by a BBSRC DTP (BB/M011186/1). DJI was supported by a National Health and Medical Research Council (NHMRC) Emerging Leadership Fellowship (GNT1195210). BPH was supported by NHMRC Leadership Fellowship (GNT1196103). KSB and EVR are affiliated to the National Institute for Health Research Health Protection Research Unit (NIHR HPRU) in Gastrointestinal Infections at University of Liverpool in partnership with the United Kingdom Health Security Agency, in collaboration with University of Warwick. The views expressed are those of the author(s) and not necessarily those of the NHS, the NIHR, the Department of Health and Social Care or UKHSA. MAC is affiliated to the National Institute for Health Research Health Protection Research Unit (NIHR HPRU) (NIHR200892) in Genomics and Enabling Data at University of Warwick in partnership with the UK Health Security Agency (UKHSA), in collaboration with Universities of Cambridge and Oxford. MAC is based at UKHSA. The views expressed are those of the author(s) and not necessarily those of the NIHR, the Department of Health and Social Care or the UK Health Security Agency.

The funders of the study had no role in study design, data collection, data analysis, data interpretation, or writing of the report.

No payment was received by any pharmaceutical company or other agency to write this article.

The authors were not precluded from accessing data in the study, and accept responsibility to submit for publication.

**Research in context:** *Evidence before this study:* *Salmonella* Panama has consistently been reported as a frequently isolated *Salmonella* serovar in national surveillance datasets, causing both sporadic cases and larger outbreaks. However, this picture has been masked due to most of the focus being placed on the top two serovars associated with nontyphoidal *Salmonella* (NTS) and invasive NTS disease: Typhimurium and Enteritidis. Previous works on *S.* Panama have determined transmission to include human faeces and breast milk, as well as non-human sources such as environmental reservoirs and animals (reptiles and pigs used for food). In contrast to most of the *Salmonella* serovars causing gastroenteritis, *S.* Panama has also been associated with a range of systemic infections including septicaemia, meningitis, and osteomyelitis, mostly affecting infants. The patterns of antimicrobial resistance of this pathogen were not well known, with studies reporting a mixture of different antimicrobial resistance profiles.

*Added value of this study:* Here we conducted a large-scale study of 836 globally relevant *S.* Panama isolates to understand global population structure and disease ecology. The *S.* Panama genomes used in this study were sourced from a combination of historical collections (including the first ever isolated strain of this serovar dating from 1931), national surveillance datasets, and publicly available data. Some of these isolates had been linked to travel, allowing an inferred location for understanding population structure. Overall, this assembled *S.* Panama collection spanned a range of 88 years and covered 45 countries and regions of all inhabited continents. We applied bacterial phylodynamic approaches to determine population structure, evolutionary history, and clade-specific trends of invasiveness and antimicrobial resistance correlated with geo-temporal parameters.

*Implications of all the available evidence:* This study revealed the population structure and global ecology of *S.* Panama, indicating that multidrug resistant *S.* Panama circulates in European and Asian regions, and presents an increased number of genetic markers for extra-intestinal invasiveness. The transmission and expansion of antimicrobial-resistant *S.* Panama strains presented in this study highlight the significance of supporting control and monitoring efforts from an international perspective.

## Introduction

Nontyphoidal *Salmonella* (NTS) disease poses a significant burden to public health globally, causing approximately 153 million cases and 57,000 deaths per annum. As well as being responsible for gastroenteritis in humans, NTS can also invade normally sterile body sites resulting in bacteraemia and meningitis. There were estimated to be 535,000 cases of invasive NTS (iNTS) disease in 2017, causing 77,500 deaths worldwide^1^. In humans, the clinical presentation depends upon a combination of host immune factors and bacterial features that are specific to individual *Salmonella* pathovariants^2^. The high levels of iNTS disease caused by *Salmonella enterica* serovars Typhimurium and Enteritidis in sub-Saharan Africa have generated a significant research focus on these two serovars^1^. However, little is known about other serovars, such as *S. enterica* serovar Panama (hereafter referred to as *S.* Panama), which are also associated with iNTS disease and responsible for numerous outbreaks, as reviewed previously^3^.

Whilst the majority of cases of iNTS have been identified in sub-Saharan Africa^1,4^, *S.* Panama has a very different geographical distribution. *S.* Panama has consistently been reported as a leading cause of NTS disease in Martinique, Guadeloupe, and French Guiana and is significantly associated with causing invasive infections in children^3,5^. Although the proportion of salmonellosis cases caused by *S.* Panama is high in these locations, the prevalence of antimicrobial resistance (AMR) is reportedly low^3,5^. In contrast, other regions have seen higher levels of AMR in *S.* Panama^6^. For example, in the 1970s and 1980s *S.* Panama caused public health concern in Europe after spreading through the pork industry and causing multiple hospital outbreaks^3^. *S*. Panama maintained its ranking as one of the top 20 most frequently isolated serovars until 2017 in the European Union^7^ and is a frequent cause of invasive disease, with 7% of cases historically presenting as extraintestinal infection in England. Similarly, *S.* Panama has been an important cause of iNTS in Asia, linked with high levels of AMR. Up to 83% of domestic and imported *S*. Panama isolates in Tokyo were reported to be resistant to three or more antimicrobial classes (multidrug resistant (MDR)) by the early 2000s^3^. In Oceania, *S.* Panama was one of the top three serovars isolated between 2007 and 2016 in Queensland, Australia^8^.

This study aimed to describe the genomic epidemiology and evolutionary history of *S.* Panama in order to provide a vital baseline of understanding for this globally important serovar. We used whole genome-based approaches on historical and contemporary global genomes to describe the population structure of *S.* Panama using geographically and temporally diverse datasets. We characterised four geographically-associated *S.* Panama clades and identified regional trends in AMR. Our findings prompted a phylodynamic investigation which revealed the evolution of the serovar. Finally, we determined genetic markers of invasiveness among *S.* Panama clades.

## Methods

### Study design

The *S.* Panama collection analysed in this study (*n* = 836) were derived from three public health collections (the Unité des Bactéries pathogènes entériques (UBPE), French National Reference Center for *Escherichia coli*, *Shigella*, and *Salmonella*, Institut Pasteur (Paris, France) (*n* = 559), the Gastrointestinal bacteria reference unit, UK Health Security Agency (UKHSA, London, UK) (*n* = 147), and the Microbiological Diagnostic Unit Public Health Laboratory (MDU PHL, Melbourne, Australia) (*n* = 25) and a selection of publicly available genomes obtained through EnteroBase^9^ (http://enterobase.warwick.ac.uk) (*n* = 105). It was not possible to include all publicly available genomes in our complete analysis due to the computational power required, thus we completed a supplementary analysis of all publicly available genomes using Microreact to assess representativeness and provide context (Figure S4, Supplementary Data 1, and https://microreact.org/project/spanama-hc400-369).

A description of all the 836 isolates is available with all metadata and genome accession numbers in Table S1 and Figure S5. Metadata collected from surveillance questionnaires was used to assign and infer geographical location to each isolate. When a recent trip overseas was reported, the travel destination was considered to be the inferred isolate location^10,11^. Where travel not reported, the sampling location was taken to be the inferred location. Locations were grouped geographically into six major areas based on the United Nations classification: Africa, Asia, Europe, Latin America & the Caribbean, Northern America, and Oceania.

### Procedures

Isolates from the Institut Pasteur were whole-genome sequenced either as part of the 10,000 *Salmonella* Genomes Project^12^ (*n =* 322) using the Illumina HiSeq 4000 system or at the Institut Pasteur (*n* = 237) using the NextSeq 500 system (Illumina). The *S.* Panama isolates 61-66, 86-66, and 376-66 were sequenced on a MinION Mk1C apparatus (Oxford Nanopore Technologies). Details about DNA extraction and sequencing can be found in the Supplementary Methods.

To prepare genome sequence data for downstream analysis, reads were assembled, subjected to quality control, and annotated as previously described^13^. The genomic sequences of isolates 61-66, 86-66, and 376-66 were assembled from long and short reads, with a hybrid approach and Unicycler v0.4.8^14^. All the genome sequences are available via GenBank accession numbers listed in Supplementary Table 1, and plasmids sequences under GenBank accession numbers OR797034 (pPan376-IncN), OR797033 (pPan86), OR797031 (pPan61-IncI1), and OR797032 (pPan61-IncN).

Trimmed sequencing data was mapped against *S.* Panama ATCC7378 as reference genome (accession number CP012346) using Snippy v4.6.0 (https://github.com/tseemann/snippy) with minimum coverage of 4 and base quality of 25. QualiMap^15^ v2.0 was used to assess mapping quality and only isolates with greater than 10x genome coverage were included in downstream analysis. The mean coverage depth across all isolates was 45·13x. Snippy-core v4.6.0 was used to build a reference-based pseudogenome for each isolate which was passed through snippy-clean v4.6.0 to replace non-ACTG characters with “N”. Gubbins^16^ v2·2 was used to remove recombinant regions and invariable sites. The resultant multiple sequence alignment of reference based pseudogenomes (24,843 variant sites) was used to infer a maximum likelihood phylogeny using RAxML-NG^17^ v1.0.3 with 100 bootstrap replicates to assess support. To assign clusters, RhierBAPs^18,19^ was used specifying two cluster levels, 20 initial clusters and infinite extra rounds. A maximum likelihood tree was also constructed using RAxML-NG^17^ for pPan61-IncN (alignment length = 47,695) and pPan376-IncN (alignment length = 44,651) with isolates that had >50% coverage from C2.

To determine the evolutionary history of *S.* Panama, a chronogram was produced using BactDating^20^ v1.1. The maximum likelihood phylogenetic tree was used as an input, accounting for invariant sites. BactDating’s model comparison was used to determine the most appropriate clock model (mixed arc). Effective sample size (ESS) were as follows: for mu = 501, sigma = 501, and alpha = 435. The final model was run for 10,000,000 iterations to reach convergence. All phylogenies were visualised using the Interactive Tree of Life (iTOL)^21^ v4.2.

Genetic determinants for AMR were identified using staramr v0.5.1 (https://github.com/phac-nml/staramr) against the ResFinder^22^ and PointFinder^23^ databases with default parameters (98% identity threshold and 60/95% overlap threshold respectively). Common resistance patterns were identified and the respective contigs were extracted from the assembled genomes. To understand the genetic context of regions carrying AMR determinants, the contigs were manually inspected using Artemis^24^ v10.2. The Prokka-based annotations were confirmed and updated using BLASTx^25^ v2.10.1 to compare all coding regions against the non-redundant protein sequence database. PlasmidFinder^26^ was used to predict contigs containing plasmids, and the results were compared against contigs containing AMR determinants. Finally BLASTn^25^ v2.10.1 was used to understand the wider context of the genetic region.

The invasiveness index of each isolate was calculated using previously defined methods, using the 196 top predictor genes for measuring invasiveness of *S. enterica* provided in Wheeler *et al.* (2018)^27^. We used the same pre-trained model to compare the median invasiveness index for *S.* Panama with the invasiveness index of validation strains previously described^27^.

### Statistical analysis

The distribution of invasiveness index values for each cluster were compared using the Kruskall-Wallis test implemented through GraphPad Prism v10.1. SRST2^46^ v0.2.0 was used to identify variations between the 196 genes in study isolates.

## Results

To create a relevant dataset, we sequenced the genomes of 552 *S.* Panama isolates, and included an additional 284 *S.* Panama genomes from a combination of historical collections, national surveillance datasets, and publicly available data, including the first isolate obtained in Panama in 1931 during a food poisoning outbreak in US soldiers stationed in the Canal zone (Figure 1, Table S1, and Figure S5). The majority (67% *n =* 559) of the isolates were sourced from the Institut Pasteur, Paris, France.

**Figure 1.**
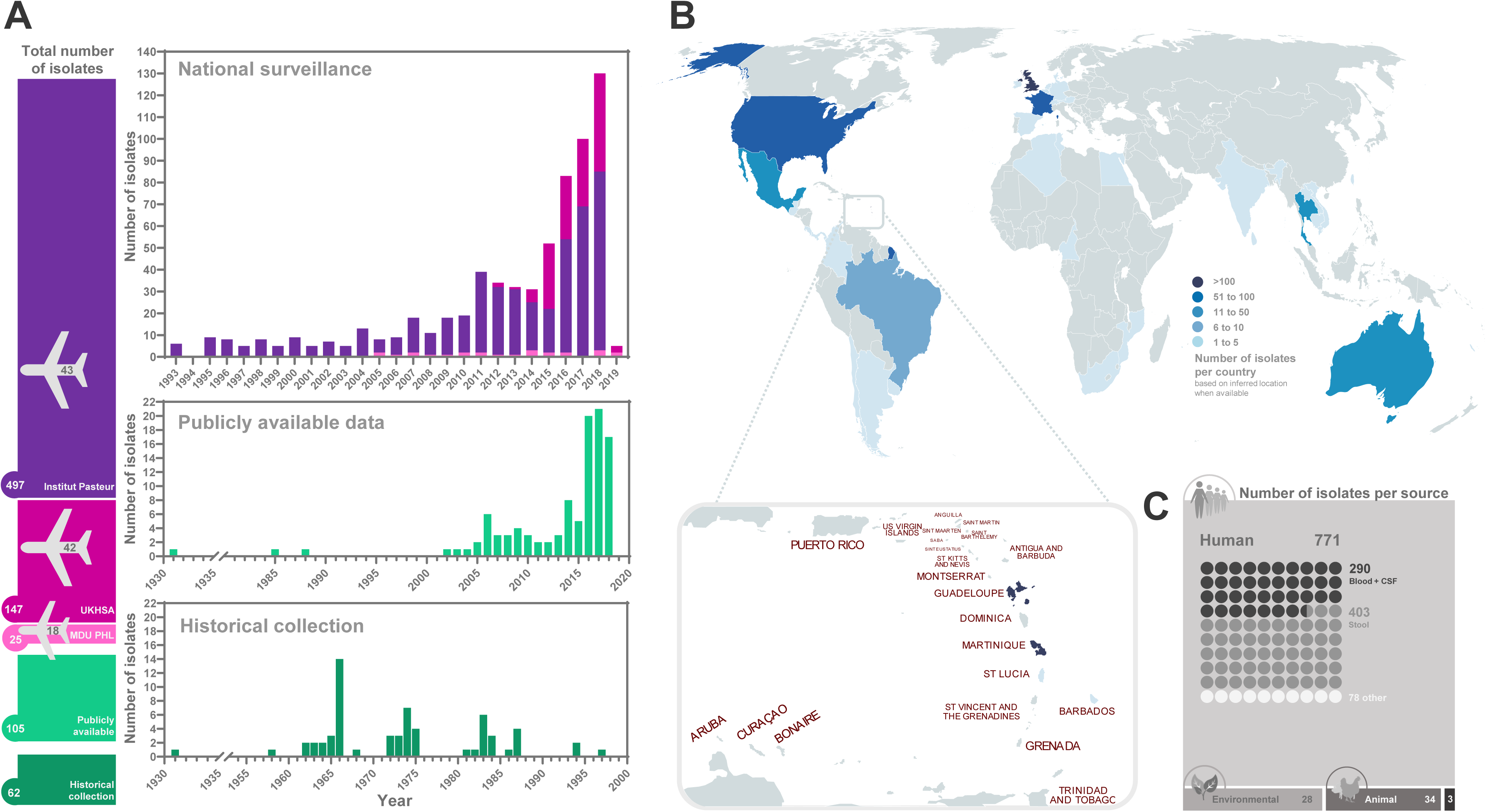
The sample collection used to construct the *S.* Panama population structure. (A) Total number of isolates per collection. Stacked colour bars on the left are proportional to the number of isolates from that source, and colours act as a key for histograms to indicate source. Airplane symbols show the number of travel-associated isolates obtained by the Institut Pasteur and United Kingdom Health Security Agency (UKHSA). Three stacked histograms show the number of isolates from UKHSA, Institut Pasteur, and MDU PHL (upper graph), publicly available data (middle graph), and historical collections (lower graph). (B) Number of isolates per region (ambiguous or outdated locations registered as “Caribbean”, “Asia”, “Antilles”, and “Yugoslavia” are omitted from the map), with regions shaded according to the inlaid key. Map created with MapChart (http://mapchart.net). (C) Number of isolates categorised by source and, for human isolates, specimen type. Three isolates of unknown source are indicated in grey. Shape sizes are proportional to the number of isolates.

For public health datasets, an isolate was deemed to be “travel associated” if a case had reported travel up to 28 days (UKHSA), seven days (MDU PHL) or 2 months (Institut Pasteur) prior to symptom onset. The “inferred region” of each isolate was set as the travel destination where available (*n =* 95) or kept as the isolation region (*n =* 741).

Overall, the collection of 836 *S*. Panama included 771 human isolates of which 403 were isolated from stool, 290 were derived from extra-intestinal sites (blood and cerebrospinal fluid), and 78 were taken from other body sites (Figure 1, Table S1). Of the 65 non-human isolates, six were of environmental origin (mainly water sources), 22 were obtained from food, 30 were from livestock, four were from wild animals, and three were of unknown origin (Supplementary Table S1). These isolates cluster with the region of origin rather than within their source group. Further work should be considered for addressing animal and environmental reservoirs.

To define population structure, we explored the phylogenetic relationship among the 836 *S.* Panama genomes and defined population groupings using Rhier Bayesian Analysis of Population Structure (RhierBAPS)^18,19^ (Figure 2). The BAPS groupings reflected four monophyletic clades with greater than 95% bootstrap support containing variable numbers of isolates. We defined the four clades as C1 (*n* = 338), C2 (*n* = 124), C3 (*n* = 131), and C4 (*n* = 104). The remaining 139 isolates were not assigned to a clade due to the level of diversity. Temporal regression of the phylogeny showed that while the inferred substitution rates were comparable among Clades C1 – C3, Clade C4 evolved nearly an order of magnitude faster (Figure S6, Table S2).

**Figure 2.**
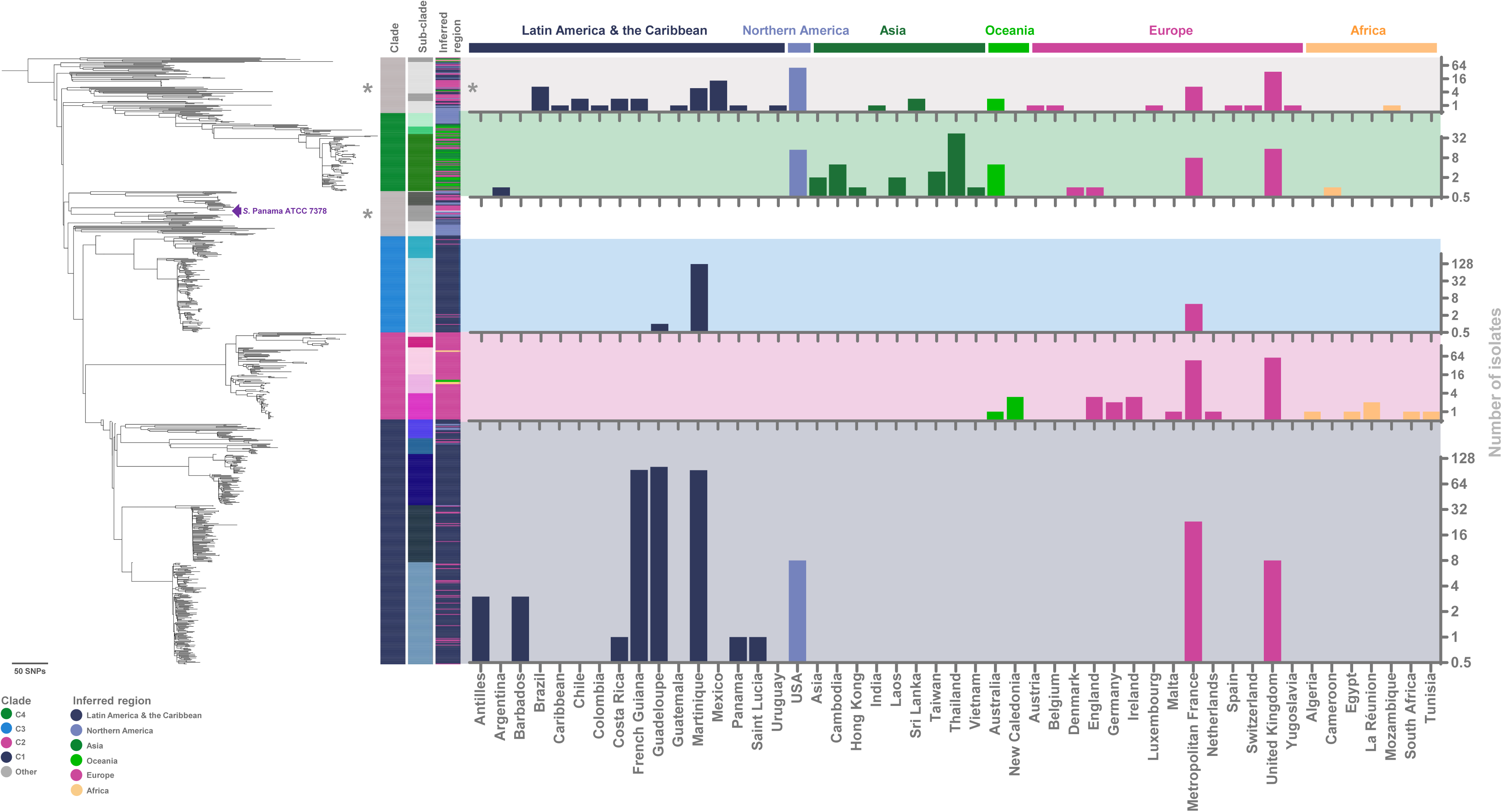
Population structure and geographical distribution of *S.* Panama isolates. Maximum likelihood phylogenetic tree of 837 isolates (including reference isolate ATCC7378 marked by a purple arrow) showing clade (first column) and subclade (second column) assignment based on RhierBAPS, inferred region (third column), and number of isolates (fourth column) as a graph with countries (x-axis) clustered and coloured by the United Nations geographical regions. Data for isolates not assigned to a clade due to the level of diversity were labelled as “other” (asterisked) and were combined into a single (asterisked) histogram marked in grey.

To investigate the relationship between clades and geographic region, location data was correlated with phylogenetic position (Figure 2). Where information on travel was available, we used the destination of travel as an inferred location. All the travel-associated isolates came from the Institut Pasteur’s national surveillance dataset (*n =* 43), UKHSA (*n =* 42), and MDU PHL (*n* = 18), leading to the inclusion of a further 12 countries not previously represented by isolates.

The majority (*n =* 295/338) of genomes clustering in clade C1 were from Latin America & the Caribbean, including the first *S.* Panama isolate from 1931. Within Clade C1, there were five geographically-linked monophyletic clusters: three were dominated by isolates from Guadeloupe, Martinique, and French Guiana respectively, and two contained isolates from multiple geographical locations in Latin America & the Caribbean (Figure 2). Clade C1 was named the “Latin America & the Caribbean clade” reflecting its geographical composition. A further clade, Clade C3 consisted of isolates from Martinique (*n =* 125/131), and was named the “Martinique clade”.

Clade C2 was mainly composed of isolates obtained from Europe, and included distinct subclades in which isolates from metropolitan France and the UK were overrepresented. Specifically, while the majority (84% *n =* 107/124) of isolates in each of these subclades came from the UK (*n =* 60/124) and metropolitan France (*n =* 47/124), C2 also contained isolates from less well sampled European countries including Germany, Malta, Ireland, and the Netherlands. These findings are consistent with European-wide circulation, and so C2 was designated as the “European clade”.

Clade C4 included isolates mainly from Asia and Oceania (*n =* 63/104). We identified three C4 subclades, of which two were dominated by isolates obtained from Asia and Oceania and a third comprised of isolates obtained in the USA. Accordingly, C4 was termed the “Asia/Oceania clade”.

The majority of genomes obtained from travel associated cases were related to travel to Asia (*n* = 52) and Latin America & the Caribbean (*n* = 36), which mainly cluster within C4 (Asia) and C1 & C3 (Latin America & the Caribbean). To assess representativeness, we placed our collection into the context of all 3,051 publicly available *S*. Panama genomes (using core genome multilocus sequence typing hierarchical cluster HC400 = 369, as of October 2024) in EnteroBase^9^ and completed a supplementary analysis using Microreact (Figure S4, Supplementary Data 1, and https://microreact.org/project/spanama-hc400-369). This comparison suggests that our collection provides good representation of publicly accessible data.

Of the 836 isolates, 14% (*n =* 121/836) were predicted bioinformatically to be resistant to at least one antimicrobial class. Previous studies have demonstrated that genome-based AMR analysis accurately predicted phenotype for 98% of *Salmonella* isolates^28^. The remaining 86% (*n* = 715/836) isolates were predicted to be pan-susceptible, lacking any known antibiotic resistance genes or mutations. Most resistant isolates in our collection (*n =* 113/121) fell within C2 (European clade) and C4 (Asia/Oceania clade) (Figure 3). A search of publicly available *S.* Panama genomes revealed a small number of MDR isolates sourced in the US. The majority of the MDR US isolates were phylogenetically situated with isolates in our collection that belonged to C4 (Asia/Oceania clade) or had clade unassigned (Supplementary Figure S4, Supplementary Data 1, and https://microreact.org/project/spanama-hc400-369).

**Figure 3.**
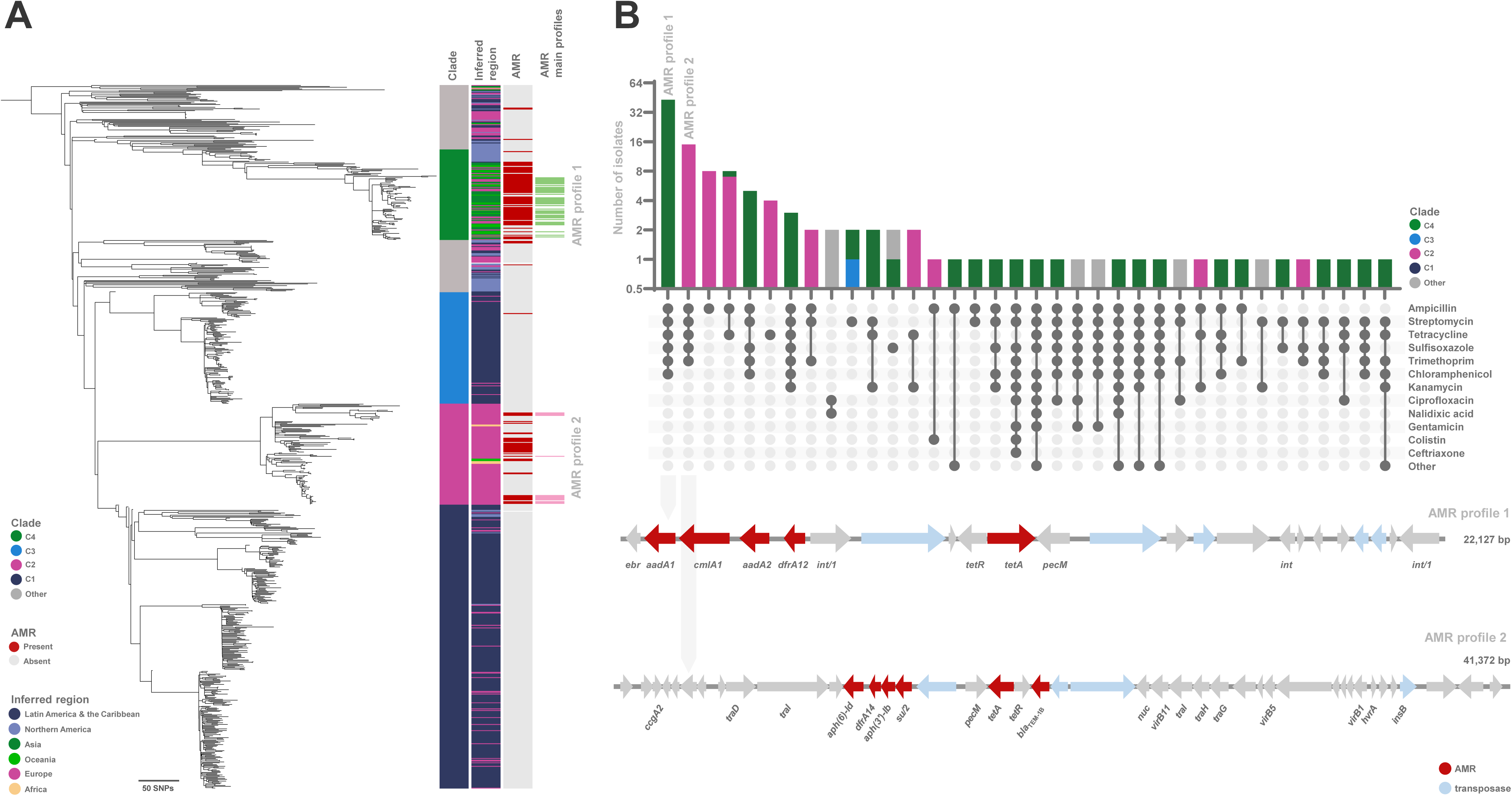
Antimicrobial resistance trends in *S.* Panama. (A) Maximum likelihood phylogenetic tree showing clades (first column), inferred region (second column) coloured based on UN geographical regions, AMR (third column) presence of at least one resistance conferring gene (red) or absence (grey), and presence of predominant AMR profiles: AMR profile 1 (green) and profile 2 (pink). (B) Number of isolates (coloured by clade) with different AMR profiles. Resistance to different antimicrobials is shown as present (dark circles) or absent (light circles). Dark circles are connected to highlight a variety of unique AMR profiles. The two predominant AMR profiles (AMR profile 1: resistance to ampicillin, streptomycin, tetracycline, trimethoprim, and chloramphenicol, and AMR profile 2: resistance to ampicillin, streptomycin, tetracycline, trimethoprim, and kanamycin) are highlighted. Functional annotation of contiguous sequences with the two predominant AMR profiles (respectively indicated by grey arrows pointing to sequence schematics). Arrows indicate predicted reading frames highlighting genes encoding AMR (red) and transposases (light blue).

The most common resistance profile, occurring in 36% (*n =* 43/121) of resistant isolates, was against streptomycin (*aadA1, aadA2*), ampicillin (*bla*_TEM-1B_), chloramphenicol (*cmlA1*), trimethoprim (*dfrA12*), sulfisoxazole (*sul3*), and tetracycline (*tetA*). Detailed analysis of the draft genomes showed that most of the resistance genes (*aadA1*, *aadA2*, *cmlA1*, *dfrA12* and *tetA*) were encoded by an MDR cassette located on a single contiguous sequence (contig; Figure 3B). A BLAST search of the contig revealed 99·8% sequence identity to a 21·2 kb region of an 83·2 kb plasmid (GenBank accession number CP044301) which had previously been identified in an *E. coli* isolate obtained from pork in Cambodia^29^.

The second most common AMR profile, found in 12% (15/121) of resistant isolates was against streptomycin (*aph(3’’)-Ib* and *aph(6)-Id*), ampicillin (*bla*_TEM-1B_), trimethoprim (*dfrA14*), sulfisoxazole (*sul2*), and tetracycline (*tetA*). All genes were carried by an MDR cassette located on a single contig (Figure 3B). The contig shared 99·9% sequence identity with a 28·3 kb region of a 50·9 kb *S. enterica* plasmid (GenBank accession number CP028173), which has been identified in multiple strains including an *S.* Enteritidis isolate from Ghana^30^.

The remaining resistant *S.* Panama isolates had a variety of susceptibility profiles, including one extensively drug resistant (XDR) isolate 201209115 (GenBank accession number SAMEA6142076), which was phylogenetically situated within C4 (Asia/Oceania clade) and was isolated in 2012 from the bloodstream of a case who had reported recent travel to Thailand. The nomenclature used to describe the XDR isolate is in line with previous definitions of resistance to three first-line drugs (chloramphenicol, ampicillin, and trimethoprim-sulfamethoxazole) as well as fluoroquinolones and third-generation cephalosporins^31^. The isolate carried AMR determinants against eight different antimicrobial classes, including: fluoroquinolones, polymyxins, and extended spectrum beta-lactamase (ESBL) mediated resistance to third generation cephalosporins (*bla*_CTX-M-55_). The resistance genes in the XDR isolate were distributed across five contigs (Figure S1), with the kanamycin (*aph(3’)-Ia*) and gentamicin (*aac(3)-IId*) genes being carried on separate contigs. Genes encoding resistance to streptomycin (*aadA1, aadA2*), chloramphenicol (*cmlA1*), trimethoprim (*dfrA12*), and sulfisoxazole (*sul3*) were co-located on the same contig, and the resistance genes against ciprofloxacin (*qnrS1*), ampicillin, and ceftriaxone (*bla*_CTX-M-55_) were also co-located on a separate contig.

Due to resource constraints, it was not possible to conduct a contig-based analysis for all isolates, consequently a short-read based analysis was initially conducted followed by targeted resequencing with Oxford Nanopore. Our short read-based plasmid prediction analysis identified that only 16% (*n =* 131/836) of *S.* Panama isolates in this study carried plasmids, including 65% (*n =* 85/131) of resistant isolates and 35% (*n =* 46/131) susceptible isolates. The main plasmid incompatibility groups identified were IncFIA(HI1) (*n* = 40/131) and IncFIB(K) (*n* = 35/131) restricted to C4 (Asia/Oceania clade), with the main AMR profile streptomycin, ampicillin, chloramphenicol, trimethoprim, sulfisoxazole, and tetracycline (*n* = 27/35). IncN plasmids were identified mostly in C2 (European clade) (*n* = 28/131) that shared the same AMR profile as IncFIA(HI1) and IncFIB(K) plasmids, but includes kanamycin resistance instead of chloramphenicol resistance.

To explore AMR context and evolution within C2 (which shows high levels of resistance), plasmid DNA of three MDR isolates (isolates 61-66, 86-66, and 376-66) were extracted and sequenced using long-read sequencing technology (Oxford Nanopore). These isolates were collected during hospital outbreaks caused by *S*. Panama in France in 1966 and clustered within C2 (European clade). Sequence analysis showed a 45 kb IncN plasmid (pPan376-IncN) encoding resistance to tetracycline (*tetA*), and a small 8 kb plasmid (pPan86) encoding resistance to ampicillin (*bla*_TEM-1_). Functional annotation of the plasmids identified in isolate 61-66 showed a 104 kb IncI1 plasmid (pPan61-IncI1) with integration of the *bla*_TEM-1_ gene, and a 48 kb IncN plasmid (pPan61-IncN) encoding tetracycline and ampicillin resistance, potentially from the fusion of pPan376-IncN and pPan86 (Figure S2).

To understand whether these IncN plasmids are driving resistance in recent European clade C2 isolates, we screened for the presence of plasmids in the genomes of all C2 isolates. This screening indicated that 23% (*n* = 27/124) of isolates contained pPan376-IncN, 26% (n = 32/124) contained pPan86, 2% (*n* = 3/124) pPan61-IncI1, and 23% (*n* = 28/124) pPan61-IncN (Figure S2 and Table S1). We constructed phylogenies of the two most prevalent IncN plasmids to explore coevolution and determine whether these plasmids are continuing to drive current AMR in *S.* Panama (Figure S2). The results show temporal clustering, indicating that the older and contemporary IncN plasmids are distinct.

To investigate the evolutionary dynamics of the global *S*. Panama population, we inferred the times to most recent common ancestor (MRCA) of each clade using BactDating (Figure 4)^20^. Analysis of the chronograph predicted the MRCA of all isolates in this study to be year 1555 with a highest probability density interval (HPD) of 1458 to 1634. C1 (Latin America & the Caribbean clade) had an MRCA of 1899 (HPD 1880-1911) with separate introductions into Guadeloupe, Martinique, and French Guiana, followed by clonal expansion in the 1930s and 1950s in each of the three regions. The MRCA of C2 (European clade) was 1879 (HPD 1852-1894), followed by expansion of more contemporary subclades in the 1970s. The MRCA of C3 (Martinique clade) is 1880 (HPD 1862-1916), and of C4 (Asia/Oceania clade) is 1879 (HPD 1854-1905). These predicted timeframes overlap with European attempts to, and the final construction of, the Panama Canal.

**Figure 4.**
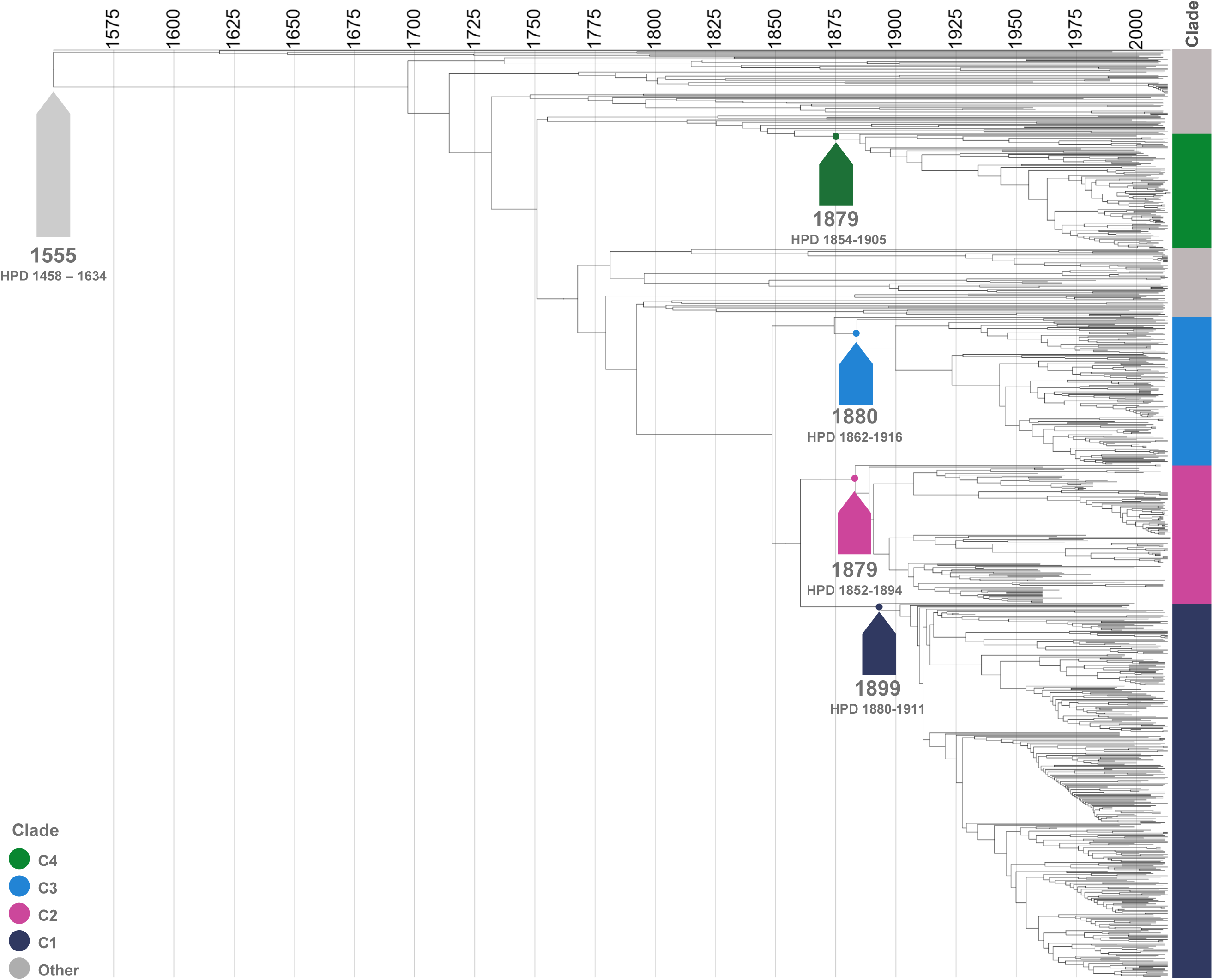
Phylodynamic and evolutionary timescales of *S.* Panama. Chronograph for estimation of most recent common ancestors for *S*. Panama clades. Arrows indicate MCRA for each clade, with year and highest probability density (95%) interval shown as text.

Whilst it was beyond the scope of this study to provide a detailed analysis of all possible markers of invasiveness, we provide a snapshot of the invasive potential of *S.* Panama using the “invasiveness index” model^27^ (Table S1). The model provides a prediction of invasiveness by assessing genome degradation within 196 genes and has been validated for other *Salmonella* serovars^27^. The mean invasiveness index of *S.* Panama was 0·2294 (SD = 0·01790). Within *S.* Panama, C2 had the greatest mean invasiveness index (European clade; median (IQR) was 0·2474 (0·2336-0·2502), followed by C3 (Martinique clade; median (IQR) was 0·2343 (0·2229-0·2367), C1 (Latin America & the Caribbean clade; median (IQR) was 0·2303 (0·2210-0·2408), and C4 (Asia/Oceania clade; median (IQR) was 0·2283 (0·2186-0·2407)).

## Discussion

This study has described the genomic epidemiology and evolutionary history of *S.* Panama, providing a vital baseline of understanding for this globally important serovar with relevance to genome-based surveillance internationally. Our finding revealed the presence of four geographically-linked phylogenetic clades, with C1 consisting predominantly of isolates from Latin America & the Caribbean, C2 from Europe, C3 from Martinique and C4 from Asia/Oceania.

The majority of *S.* Panama in this study were pan-susceptible to antibiotics, however there are a number of MDR isolates in C2 (European clade) and C4 (Asia/Oceania clade) (Figure 3), of which the former presented genomic predictors of elevated invasiveness (Figure S3B). These findings are supported by reports of an increase in *S.* Panama isolates from Asia that were resistant to multiple antibiotics, including cotrimoxazole, ampicillin, streptomycin, kanamycin, and gentamicin^32^.

We also identified clustered AMR elements on the genome that are consistent with plasmid-mediated resistance in *S.* Panama, as has been seen in previous studies^32^. The plasmid complement of the *S*. Panama serovar has rarely been studied in the past, but it has been reported that *S.* Panama does not commonly carry the large virulence plasmids that have previously been characterised in other *Salmonella* serovars^33^. Rather, *S.* Panama have previously been found to carry a heterogeneous population of plasmids (Table S1)^34^.

The evolutionary dynamics analysis of the global *S*. Panama population provided information about the MRCA of each clade (Figure 4). The four clades were predicted to share a most recent common ancestor in the 1500s and have undergone contemporary introductions into different geographical regions. We identified isolates from both Guadeloupe and Martinique in C1 and C3, suggesting that inter-island transmission of C1 is likely occurring. We speculate that C2 arose from an outbreak that involved the continental spread of *S.* Panama across Europe, associated with the food production industry in the 1970s and 1980s^3^. It is possible that these European *S*. Panama were exposed to significant levels of antibiotics in humans and/or food animals, which selected for resistance to multiple antibiotic agents via acquisition of mobile genetic elements^3^, consistent with the indicators of mobilisation (Figure 3). The emergence of C2 (European clade) in the 1970s is consistent with this hypothesis, coinciding with the propagation of *S.* Panama in Europe via the pig industry^3^. To note, we identified an MDR isolate from swine in the US (SRR8381713) as part of our analysis of all publicly available data. However, this isolate was phylogenetically situated in C4 (Asia/Oceania clade).

Although *S.* Panama causes extraintestinal disease globally, genetic determinants of invasiveness have never been investigated. In this context, invasiveness reflects the ability of the pathogen to cause systemic spread and extraintestinal infection in humans^35^. Invasive infections cause much higher levels of mortality than gastroenteritis; for example, the case-fatality rate of iNTS is 20%^1^, whereas the case-fatality rate of *S.* Typhimurium-associated gastroenteritis is 0·6%^36^. It has been observed that functions required for escalating growth within the inflamed gut are often lost among iNTS variants in a series of gene degradation/pseudogenisation events^13^. The increased clinical invasiveness observed amongst *S.* Panama is likely influenced by several factors, related to both the host (e.g. immunocompetency) and pathogen (e.g. presence/absence of genes that elevate invasiveness). The calculated invasiveness index of *S.* Panama is slightly higher than that described for serovars with a typically broader host range (Agona, Enteritidis, Heidelberg, Newport, and Typhimurium)^27^ but significantly lower than that of host-restricted serovars (Typhi, Paratyphi, Gallinarum, Pullorum, Dublin, and Choleraesuis)^27^ (Figure S3A). Public health surveillance should take into consideration the significantly higher invasiveness index of C2 (European clade) combined with the high proportion of AMR isolates when risk assessing outbreaks falling within this clade (Figure S3B).

There are limitations associated with travel-related data in this study. For example, it is possible that cases who reported travel did not necessarily acquire their infection whilst travelling. Furthermore, information on travel was only available for public health datasets with limited (likely under-reported) or unavailable travel information for all cases. Thus, there are likely some geographical signals missing which may impact the interpretation of geographically-linked clade designations. However, we note that for all four clades in which a geographical designation has been made, there are a substantial number of isolates from the relevant region. This is indicative of domestic circulation, but does not totally exclude the possibility that isolates were initially imported via travel. Moreover, our analysis only shows the population structure of those isolates selected for inclusion in this *S.* Panama collection. Thus, the analysis may be subject to a number of sampling biases. Isolates submitted to national surveillance teams at public health centres will be limited to those sampled from cases who attend healthcare services and have a sample collected, which may favour more clinically severe cases. Also, not all isolates submitted to national surveillance will be whole-genome sequenced and uploaded to publicly available repositories, with approaches differing by country. The publicly available datasets may also include isolates submitted as part of research projects, each of which will have had its own sampling strategy that may have favoured sequencing of isolates with specific characteristics, for example, those with specific antimicrobial resistance patterns. Finally, the analysis is limited by the sampling strategy which purposively selected for temporal and geographical breadth. As a result, our analysis likely captures only a partial view of the true global population structure of *S.* Panama.

As the first large-scale phylogenetic analysis of the *S.* Panama serovar to our knowledge, we have revealed important information about population structure, AMR, global ecology, and invasiveness. It will be important to monitor the incidence of clades C2 and C4 in ongoing genome-based surveillance, and to understand the movement of mobile genetic elements responsible for antibiotic resistance. It is possible that the evolutionary trajectory of *S*. Panama will parallel the *Salmonella* serovars Enteritidis and Typhimurium, which contain specific lineages and pathovariants with an increased invasiveness index and are now causing significant levels of human bloodstream infections^1,27,37^. Future work on *S.* Panama can be derived from the current knowledge of serovars Enteritidis and Typhimurium and should focus on understanding the molecular basis of systemic invasion, the development of a robust typing system, and a defined animal infection model.

## Acknowledgments

We thank all the corresponding laboratories of the French National Reference Center for *Escherichia coli*, *Shigella*, and *Salmonella*. We thank Dr Edward Cunningham-Oakes and Dr Nicolas Wenner for useful discussions.

## Appendix

### Supplementary Methods

**Supplementary Data 1: Microreact project of All S. enterica HC400 = 369 available in EnteroBase**

**Table S1: S. Panama metadata and accession numbers**

**Table S2: Nucleotide substitution analysis of four S. Panama clades**

**Figure S1: Functional annotation of five AMR gene cassettes in XDR isolate 201209115**

The predicted phenotype conferred (left) and total length (right) for each contiguous sequence that carries an AMR gene cassette is displayed. Arrows indicate open reading frames, and AMR genes are highlighted in red. GenBank accession number SAMEA6142076.

**Figure S2: Conservation and functional annotation of plasmids within C2 ascertained with long-read sequencing technology**

(A) Maximum likelihood phylogenetic tree of C2 showing subclades (column one), inferred region (column two) of isolation coloured by UN geographical regions in Figure 2, year (column three), AMR (column four) presence (red) or absence (grey), and percentage coverage (>50%) of pPan61-IncI1 (column five), pPan61-IncN (column six), pPan376-IncN (column seven), and pPan86 (column eight). (B) Pairwise comparison of linearised plasmids showing genomic similarity (red) of pPan376-IncN and pPan86 with pPan61-IncN. GenBank accession numbers are shown below the plasmid name. (C) Phylogenetic trees of pPan376-IncN and pPan61-IncN showing year of isolation (first column) and AMR (following column(s)). The AMR profile of all isolates in pPan376-IncN tree was streptomycin, ampicillin, chloramphenicol, trimethoprim, sulfisoxazole, tetracycline. The sequenced plasmids that acted as reference sequences are indicated by a star. (D) Functional annotation of genes on pPan61-IncI1, pPan61-IncN, pPan376-IncN, and pPan86.

**Figure S3: Invasiveness index of *S.* Panama clades in comparison with common *S. enterica* serovars**

(A) Median invasiveness index of different *Salmonella* serovars as calculated in Wheeler et al (2018) (black dots) with *S.* Panama (this study) highlighted in pink. (B) Distribution of invasiveness index of all 836 isolates clustered by clade. Overlying box and whiskers plots indicate median and interquartile range, which are enumerated below in the table. Comparisons between groups are indicated by lines uppermost with *p*-values indicated above each line.

**Figure S4: All *S. enterica* HC400 = 369 available in EnteroBase**

Screenshot of Microreact project containing the data publicly available (October 2024) in EnteroBase (https://enterobase.warwick.ac.uk/) for all *S*. Panama genomes identified (search criteria: experimental data HC400 = 369, total of 3,051 assembled genomes) accessible in https://microreact.org/project/spanama-hc400-369 and Supplementary Data 1.

**Figure S5: *S.* Panama phylogenetic tree with annotated tips**

High resolution maximum likelihood phylogenetic tree showing clades (first column), inferred region (second column) coloured based on UN geographical regions, and AMR (third column) presence of at least one resistance conferring gene (red) or absence (grey). All genome accession numbers are annotated in tree tips, corresponding with data available in Supplementary Table S1.

**Figure S6. Nucleotide substitution analysis of four *S.* Panama clades.**

Root-to-tip distance extracted from the maximum likelihood phylogenetic tree of 837 isolates (excluding reference isolate ATCC 7378) plotted against year of isolation. Data available in Supplementary Table S2.

