## Supplementary Figures for "Global diversity and evolution of *Salmonella* Panama, an understudied serovar causing gastrointestinal and invasive disease worldwide: a genomic epidemiology study"

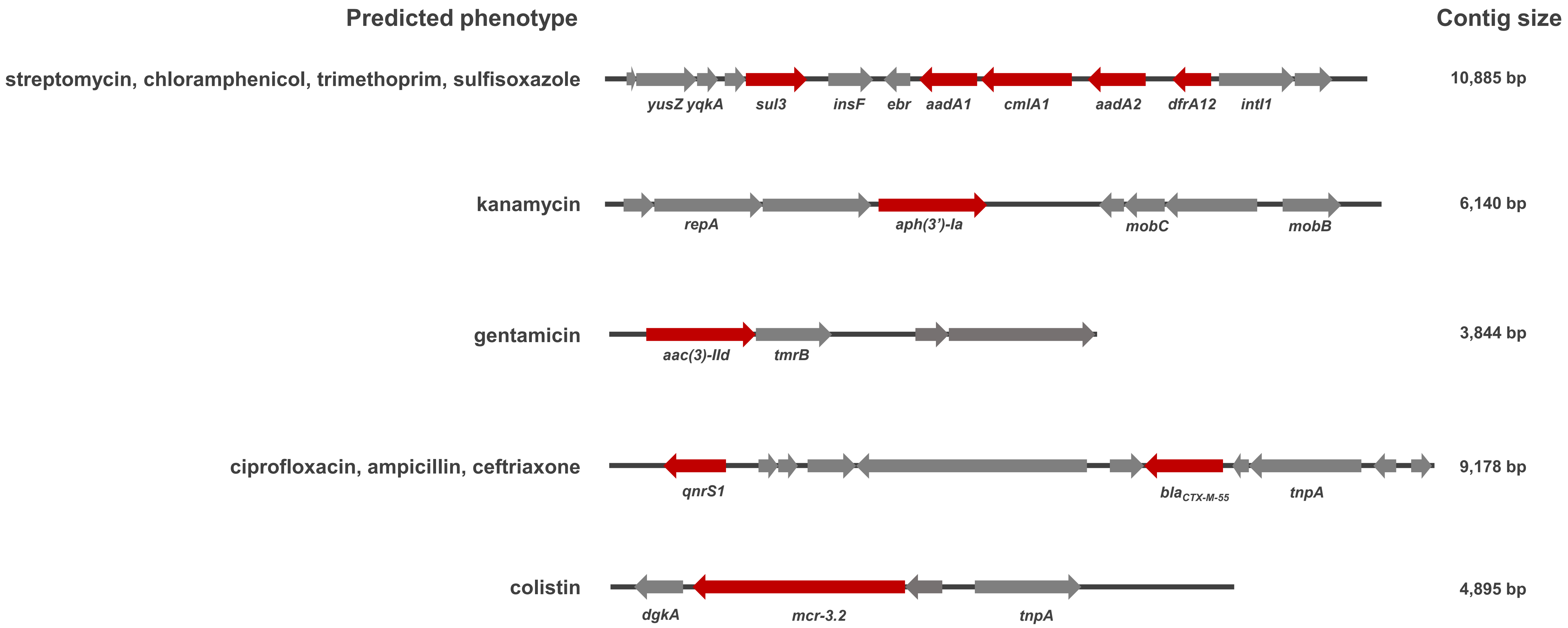

Predicted phenotype
Contig size
streptomycin, chloramphenicol, trimethoprim, sulfisoxazole
yusZ yqkA
sul3
insF
ebr
aadA1
cmlA1
aadA2
dfrA12
intI1
10,885 bp
kanamycin
repA
aph(3')-Ia
mobC
mobB
6,140 bp
gentamicin
aac(3)-IId
tmrB
3,844 bp
ciprofloxacin, ampicillin, ceftriaxone
qnrS1
blaCTX-M-55
tnpA
9,178 bp
colistin
dgkA
mcr-3.2
tnpA
4,895 bp

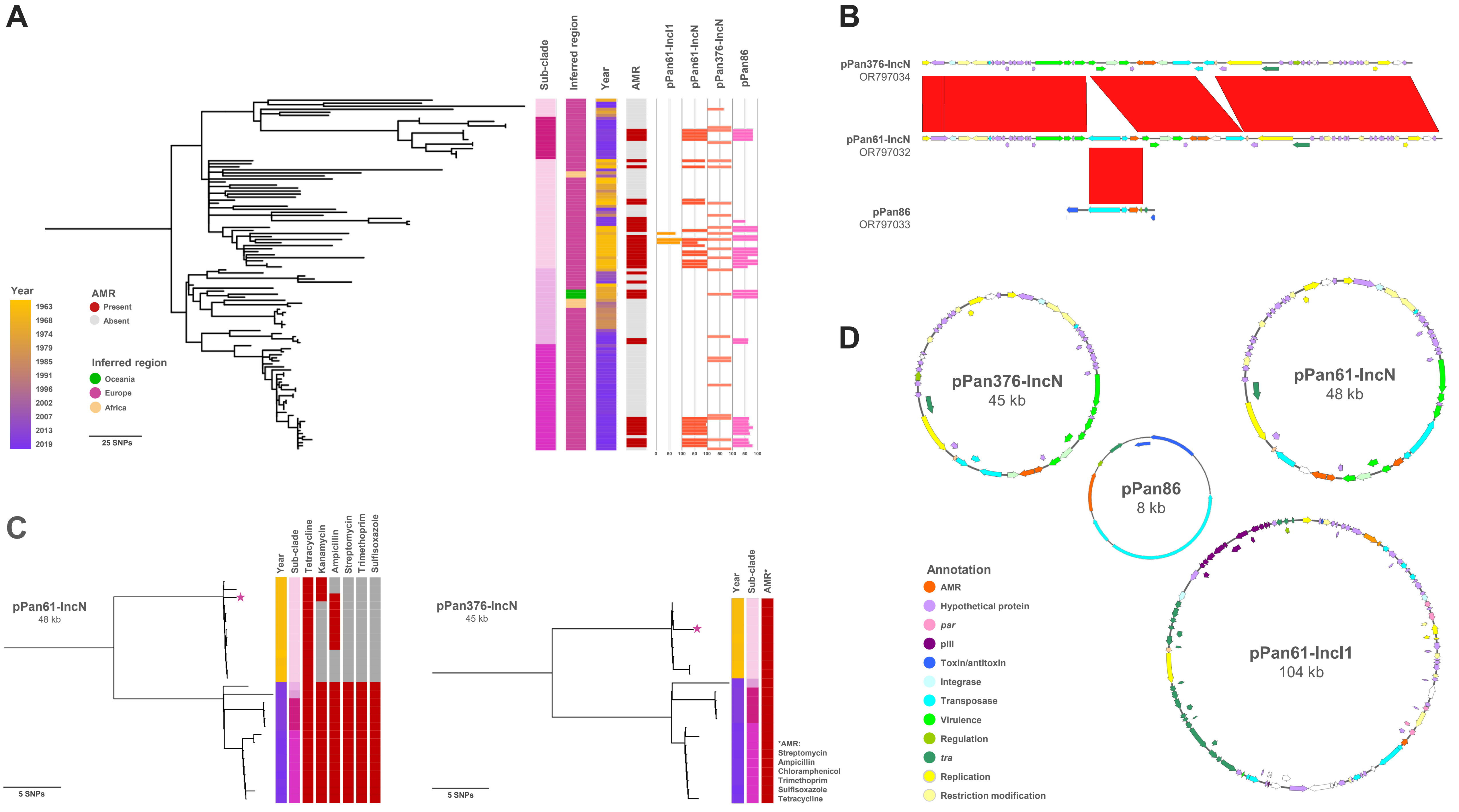
A
Sub-clade
Inferred region
Year
AMR
pPan61-IncI1
pPan61-IncN
pPan376-IncN
pPan86
Year
1963
1968
1974
1979
1985
1991
1996
2002
2007
2013
2019
AMR
Present
Absent
Inferred region
Oceania
Europe
Africa
25 SNPs
B
pPan376-IncN
OR797034
pPan61-IncN
OR797032
pPan86
OR797033
C
pPan61-IncN
48 kb
Year
Sub-clade
Tetracycline
Kanamycin
Ampicillin
Streptomycin
Trimethoprim
Sulfisoxazole
5 SNPs
pPan376-IncN
45 kb
Year
Sub-clade
AMR*
*AMR:
Streptomycin
Ampicillin
Chloramphenicol
Trimethoprim
Sulfisoxazole
Tetracycline
5 SNPs
D
pPan376-IncN
45 kb
pPan61-IncN
48 kb
pPan86
8 kb
pPan61-IncI1
104 kb
Annotation
AMR
Hypothetical protein
par
pili
Toxin/antitoxin
Integrase
Transposase
Virulence
Regulation
tra
Replication
Restriction modification

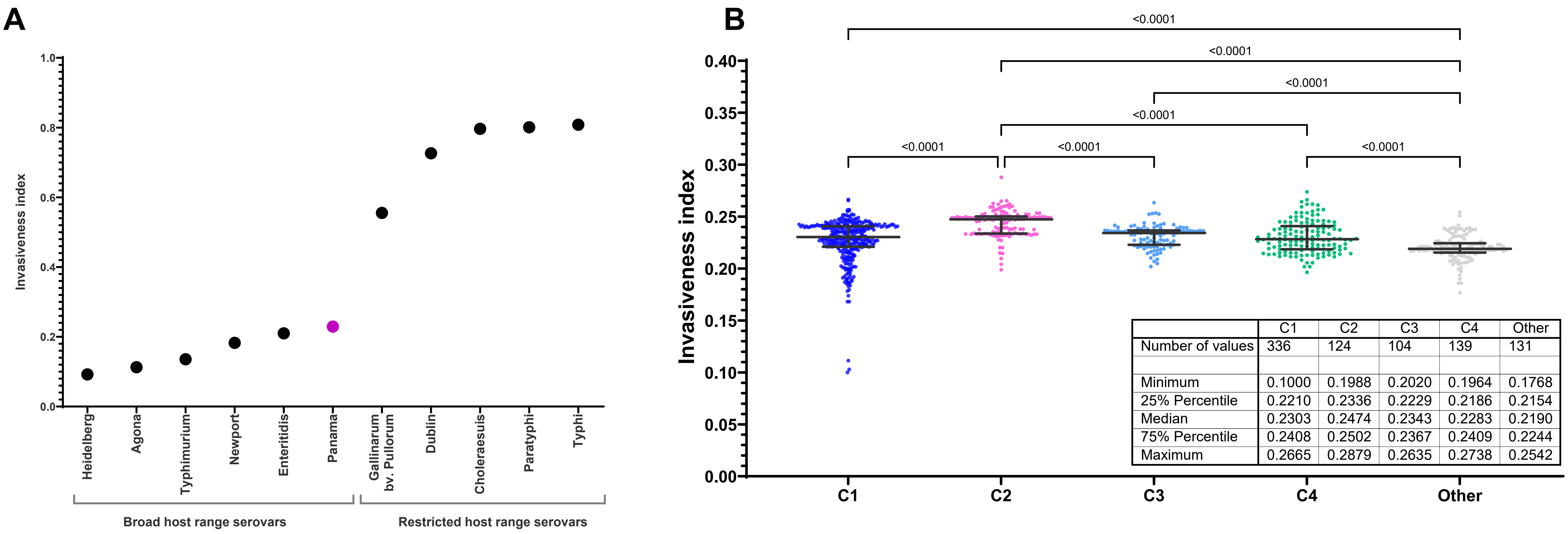
A
Invasiveness index
1.0
0.8
0.6
0.4
0.2
0.0
Heidelberg
Agona
Typhimurium
Newport
Enteritidis
Panama
Gallinarum bv. Pullorum
Dublin
Choleraesuis
Paratyphi
Typhi
Broad host range serovars
Restricted host range serovars
B
<0.0001
<0.0001
<0.0001
<0.0001
<0.0001
<0.0001
<0.0001
Invasiveness index
0.40
0.35
0.30
0.25
0.20
0.15
0.10
0.05
0.00
C1
C2
C3
C4
Other
| C1 | C2 | C3 | C4 | Other
Number of values | 336 | 124 | 104 | 139 | 131
Minimum | 0.1000 | 0.1988 | 0.2020 | 0.1964 | 0.1768
25% Percentile | 0.2210 | 0.2336 | 0.2229 | 0.2186 | 0.2154
Median | 0.2303 | 0.2474 | 0.2343 | 0.2283 | 0.2190
75% Percentile | 0.2408 | 0.2502 | 0.2367 | 0.2409 | 0.2244
Maximum | 0.2665 | 0.2879 | 0.2635 | 0.2738 | 0.2542

S. Panama in Enterobase - HC400 369

SEARCH IN ALL COLUMNS 2842 of 2842

Tree

Nodes & Labels | Metadata blocks

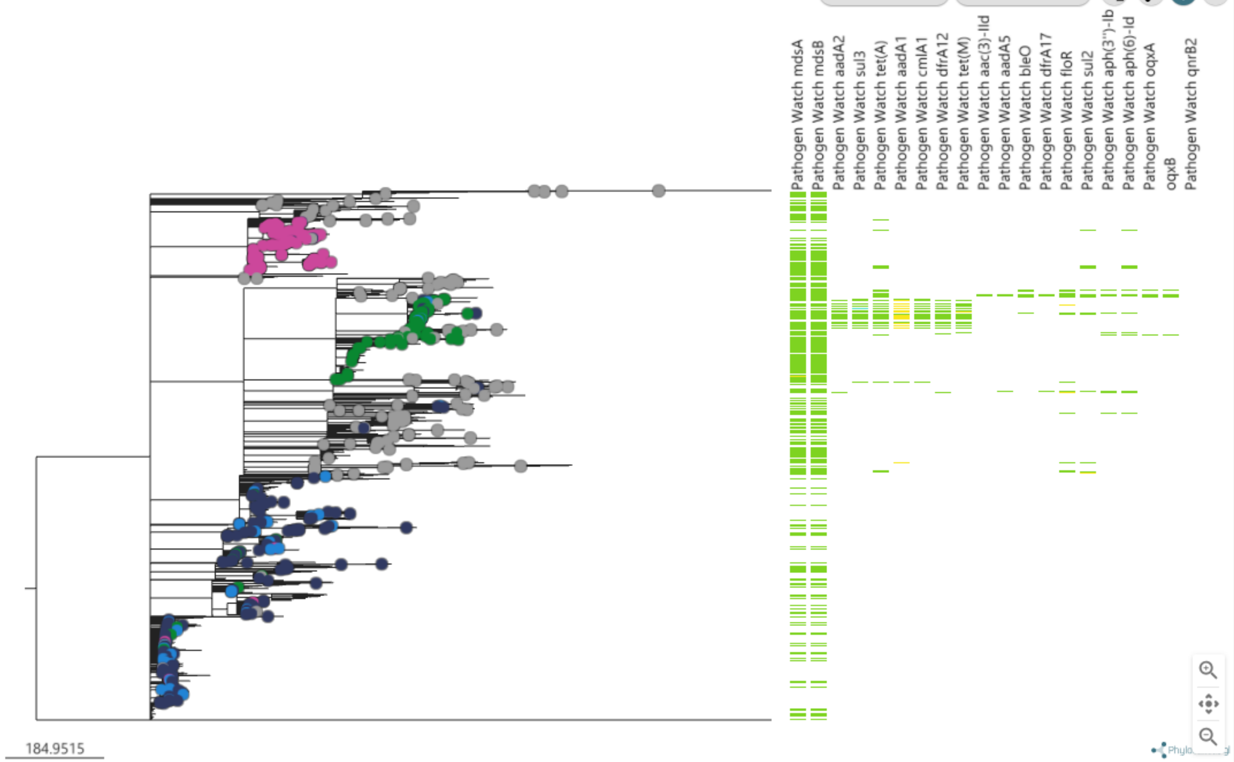


Map

Markers | Style

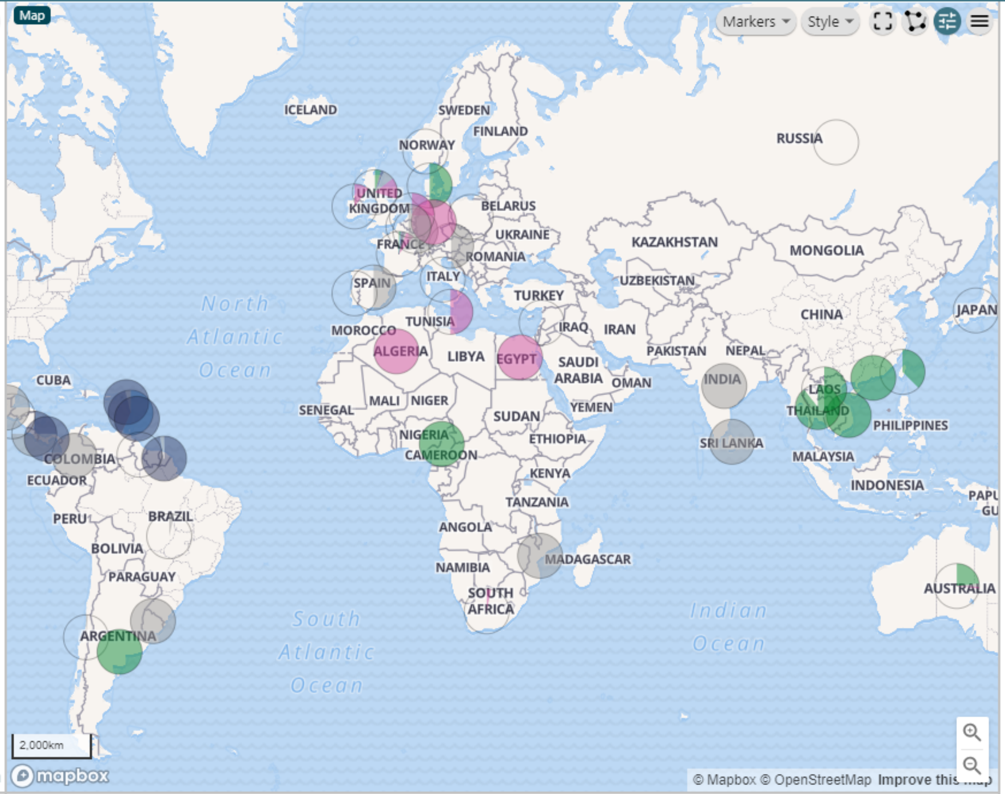


Timeline

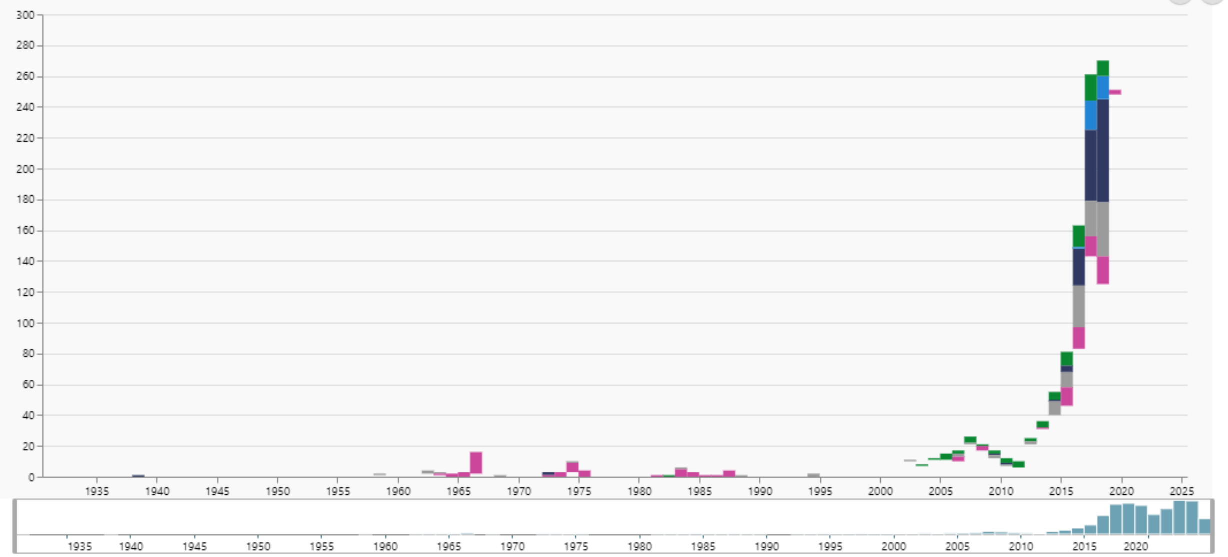


Metadata

Columns | Density

| ID | Accession No | Country | Year | Collection (Pı | Lineage (Pulf | UN Geogra |
|---|---|---|---|---|---|---|
| SAL_KC7139... | traces-0PVd... | Metropolita... | 2018 | Enterobase | | Europe |
| SAL_IB7094... | traces-0RIts... | Metropolita... | 2018 | Enterobase | | Europe |
| SAL_JB7049... | traces-0qLA... | Metropolita... | 2017 | Enterobase | | Europe |
| SAL_BC2771... | traces-0Ouf... | Metropolita... | 2019 | Enterobase | | Europe |
| SAL_BC2772... | traces-0iYjZQI | Metropolita... | 2019 | Enterobase | | Europe |
| SAL_BC2787... | traces-0Fvm... | Metropolita... | 2019 | Enterobase | | Europe |
| SAL_BC2794... | traces-0wht... | Metropolita... | 2019 | Enterobase | | Europe |
| SAL_BC2798... | traces-0vDd... | Metropolita... | 2019 | Enterobase | | Europe |
| SAL_BC2802... | traces-0OYA... | Metropolita... | 2019 | Enterobase | | Europe |
| SAL_BC2804... | traces-0IXT... | Metropolita... | 2019 | Enterobase | | Europe |
| SAL_BC2806... | traces-0Cwi... | Metropolita... | 2019 | Enterobase | | Europe |
| SAL_BC2808... | traces-0fknu... | Metropolita... | 2019 | Enterobase | | Europe |

Legend

Colours by Lineage (Pulford & Perez-Sepulveda et al. 2024)
- C1
- C2
- C3
- C4
- Other
- (blank)

Colours by Pathogen Watch mdsA
- COMPLETE
- MISTRANSLATION
- PARTIAL_END_OF_CONTIG
- (blank)

Colours by Pathogen Watch mdsB
- COMPLETE
- MISTRANSLATION
- PARTIAL_END_OF_CONTIG
- (blank)

Colours by Pathogen Watch aadA2
- COMPLETE
- PARTIAL_END_OF_CONTIG
- (blank)

Colours by Pathogen Watch sul3
- COMPLETE
- MISTRANSLATION
- PARTIAL_END_OF_CONTIG
- (blank)

Colours by Pathogen Watch tet(A)
- COMPLETE
- PARTIAL_END_OF_CONTIG
- (blank)

Colours by Pathogen Watch aadA1
- COMPLETE
- PARTIAL_END_OF_CONTIG
- (blank)

Colours by Pathogen Watch cmlA1

Legend | Selection | History | Views

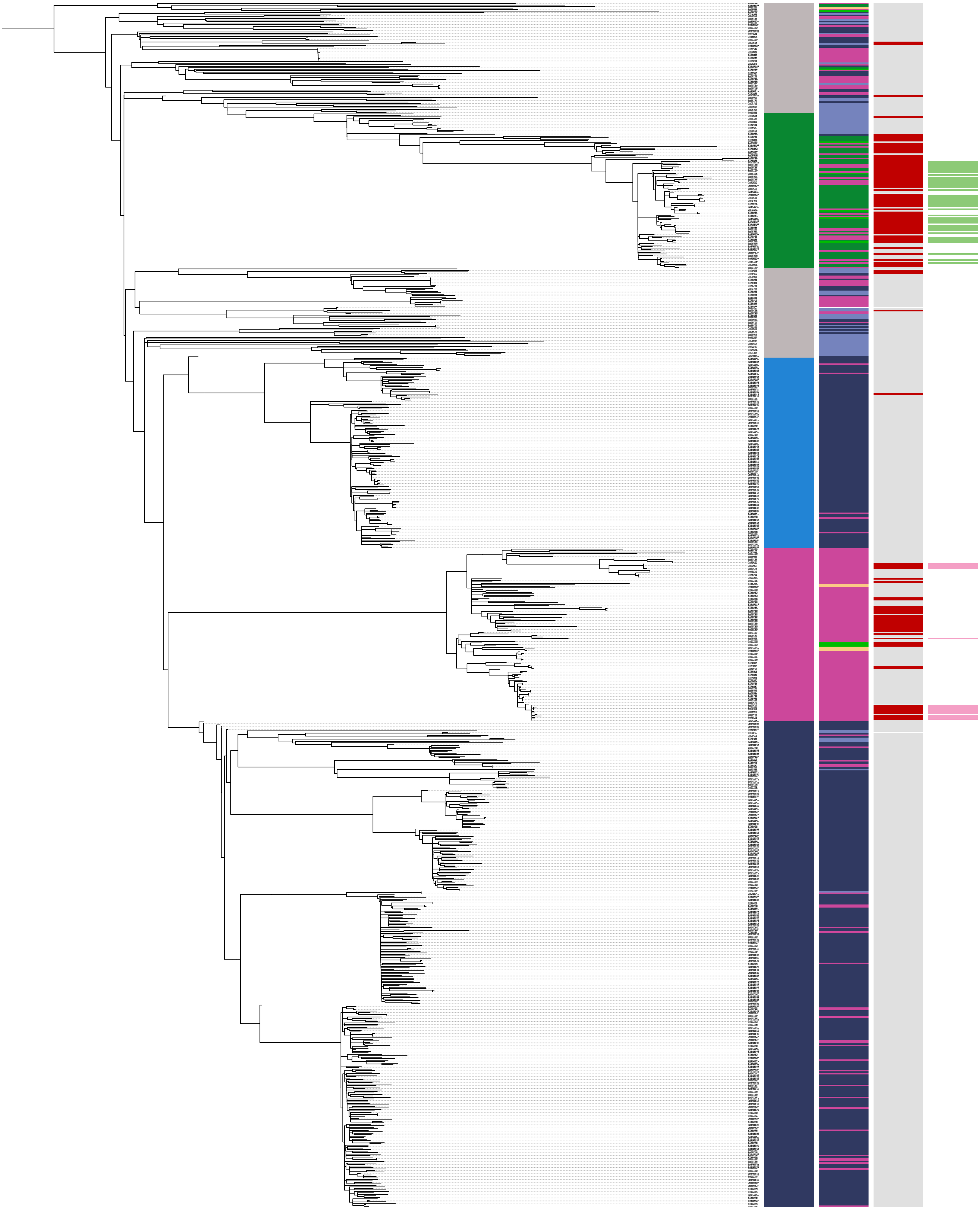

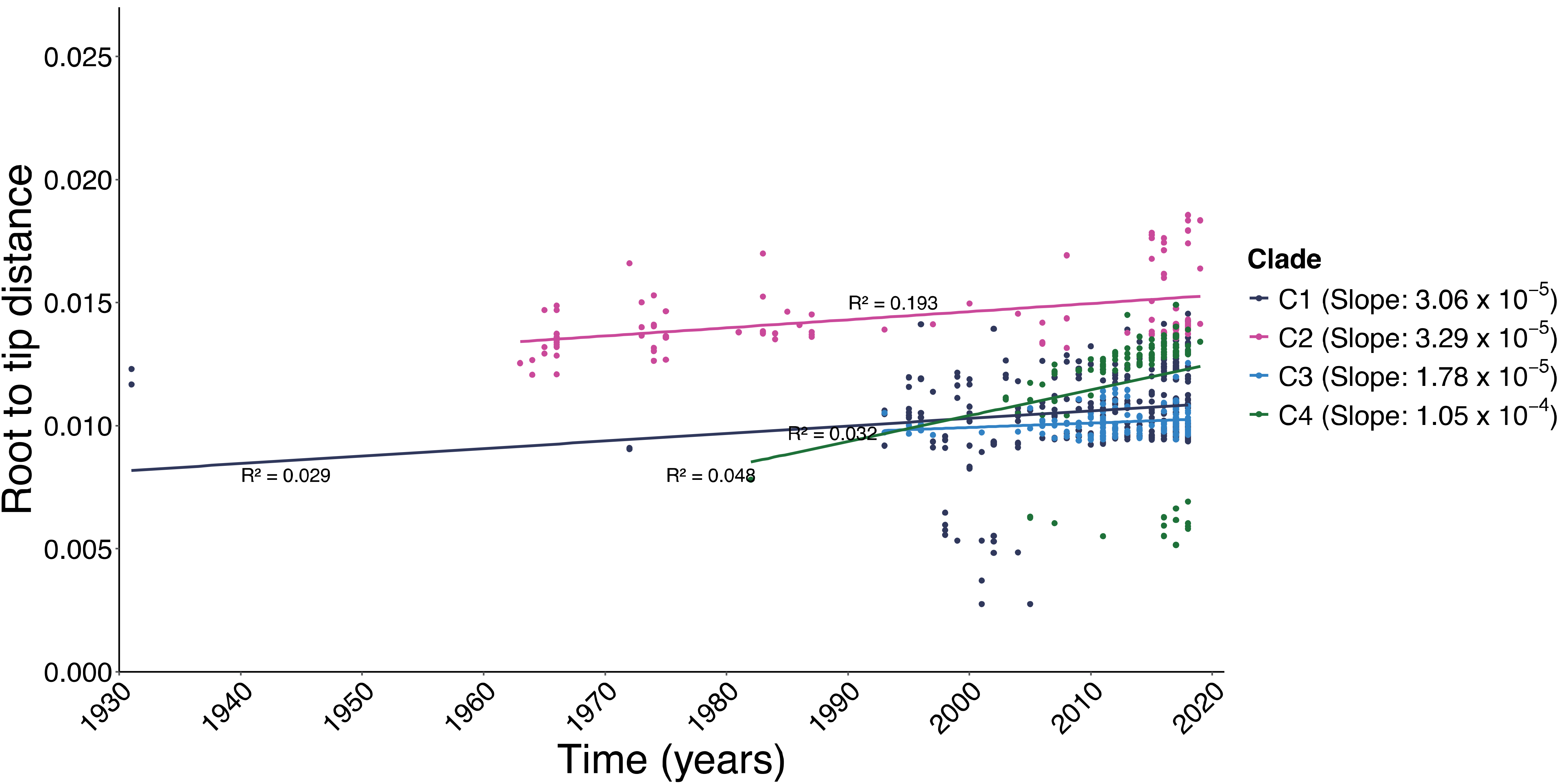

Root to tip distance
0.025
0.020
0.015
0.010
0.005
0.000
1930
1940
1950
1960
1970
1980
1990
2000
2010
2020
Time (years)
R² = 0.193
R² = 0.032
R² = 0.029
R² = 0.048
Clade
C1 (Slope: 3.06 x 10^−5)
C2 (Slope: 3.29 x 10^−5)
C3 (Slope: 1.78 x 10^−5)
C4 (Slope: 1.05 x 10^−4)
