## Supplementary Methods for "Global diversity and evolution of *Salmonella* Panama, an understudied serovar causing gastrointestinal and invasive disease worldwide: a genomic epidemiology study"

### Assembly of a *Salmonella* Panama collection with global relevance

Firstly, we selected isolates for whole genome sequencing from the Institut Pasteur UBPE’s archives which hold all *S.* Panama collected from France and French overseas territories between 1947 and 2018 (>5,000 isolates) and also reference strains, such as the 73K strain isolated in Panama in 1931. The total number of isolates that could be selected for sequencing was limited by resource constraints. We used purposive sampling of the Institut Pasteur’s collection with the aim of ensuring that the isolates selected for sequencing were as representative as possible of the temporal and global history of *S.* Panama available in the collection. Specifically, we selected 497 isolates collected for public health purposes between 1993 and 2018 (“national surveillance” isolates), and 62 isolates collected between 1931 and 1997 (“historical” isolates). This included human (*n =* 542), environmental (*n =* 10), and animal isolates (*n =* 7).

To eliminate redundancy, “national surveillance” Institut Pasteur isolates taken from the same patient prior to sampling were discarded. All isolates associated with travel outside of France and French overseas territories were selected to capture the maximum global context. All “extra-intestinal” (defined as blood and cerebrospinal fluid) isolates from French overseas territories and a small number of isolates (*n =* 90) from Metropolitan France were included. A random selection using the MS Excel random number generator tool of one stool isolate per year from each sampling location available was incorporated as non-invasive *S.* Panama isolates. Note that to eliminate regeneration of random numbers the numbers were fixed by copying and pasting as only values.

Isolates from the Institut Pasteur collection were stored as stab cultures. To resuscitate isolates, a sterile loop was used to transfer culture into 10 mL of Tryptic Soy Broth which was grown overnight at 37°C. To check for potential contaminants, each isolate was also streaked onto Drigalski agar plates, which were visually inspected following overnight growth at 37°C. If colonies were pure on Drigalski agar, 100 μL of culture was aliquoted into a FluidX 2D Sequencing Tube (FluidX Ltd, UK).

The dataset was complemented by 25 genomes generated by the Microbiological Diagnostic Unit Public Health Laboratory (MDU PHL) at The University of Melbourne, Australia, collected between 2005 and 2019^1^.

The United Kingdom Health Security Agency (UKHSA), London, UK. UKHSA (formerly Public Health England) has routinely genome-sequenced every *Salmonella* isolate received by the *Salmonella* reference service since 2015 in addition to a small number of pre-2015 isolates^2^. The resulting genomic data are regularly deposited in publicly available repositories^2^. The UKHSA dataset includes whole genome sequence data from 147 publicly available *S.* Panama collected between 2012 and 2019.

Finally, a set of 105 additional *S*. Panama publicly available genomes obtained through EnteroBase^3^ (<http://enterobase.warwick.ac.uk>) were included, which were collected between 1931 and 2018. The selection was made from 1,266 *S*. Panama genomes (*i.e.* assigned to the hierarchical cluster HC400_369 using the EnteroBase “cgMLST V2 + HierCC V1” tool^4^) on October 2020. The selection criteria considered genomes belonging to HC100 groups not previously covered and HC100 groups already described that improved the spatiotemporal or source diversity sampling. Genomes were eligible for inclusion if they had an accession number, collection year, and country of collection.

A description of all the 836 isolates is available with all metadata and genome accession numbers in Table S1 and Figure S5. A completed a supplementary analysis of all publicly available genomes from Enterobase can be found using Microreact (Figure S4, Supplementary Data 1, and <https://microreact.org/project/spanama-hc400-369>).

**Whole genome sequencing and annotation of *S.* Panama isolates**

Cultures from the Institut Pasteur stored in FluidX tubes (*n* = 315) were heat killed in a 95°C water bath for 20 min to produce thermolysates before DNA extraction (MagAttract kit, Qiagen) and whole-genome sequencing, as part of the 10,000 *Salmonella* Genomes Project^5^. Illumina Nextera XT DNA Libraries were prepared (Illumina, FC-131-1096) and sequenced (Illumina HiSeq 4000) in multiplex (768) as 150 bp paired-end reads. The rest of the isolates (*n* = 237) were grown overnight in tryptic soy broth (TSB) at 37°C, DNA was extracted using the MagNA Pure DNA isolation kit (Roche Molecular Systems, Indianapolis, IN, USA) at the Mutualized Platform for Microbiology (P2M) at Institut Pasteur, Paris. The libraries were prepared with the Nextera XT kit (Illumina, San Diego, CA, USA) and sequencing was performed with the NextSeq 500 system (Illumina) generating 150 bp paired-end reads.

Adapter content and raw read quality was assessed using FastQC v0.11.5 (<https://www.bioinformatics.babraham.ac.uk/projects/fastqc/>) and MultiQC v1.0 ([http://multiqc.info](http://multiqc.info/)). Adapter trimming was then performed using Trimmomatic^6^ v0.36 and low-quality regions were trimmed using Seqtk v1.2-r94 (<https://github.com/lh3/seqtk>) with the trimfq flag. Adapter and quality trimmed reads were assessed for quality using FastQC v0.11.5 and MultiQC v1.0. Quality metrics used to pass sequencing reads were as follows: passed basic quality statistics (total number of sequence reads, sequence length of shortest and longest read, and overall %GC), per base sequence quality, per base N content and adapter content. GC content distributions were visually inspected to determine any isolates with unusual peaks. Only high-quality reads determined using the described criteria were used in downstream analysis. Unicycler^7^ v0.3.0 was used to assemble genomes and QUAST^8^ v4.6.3 was used to evaluate assembly quality to standards consistent with Enterobase^9^. Specifically, N50 > 20 kb, 600 or fewer contiguous sequences, total number of bases between 4 and 5·8 Mbp. Prokka^10^ v1.12 was used to annotate the genomes. The *Salmonella in Silico* Typing Resource (SISTR)^11^ v1.0.2 was used to confirm serovar designations.

**Long-read sequencing and plasmid reconstruction**

The *S.* Panama isolates 61-66, 86-66, and 376-66 were cultured overnight at 37°C in alkaline nutrient agar (20 g/L casein meat peptone E2 (Organotechnie), 5 g/L sodium chloride (Sigma), and 15 g/L Bacto agar (Difco), adjusted to pH 8·4). Single colonies were used to inoculate 20 mL of Brain-Heart-Infusion (BHI) broth, and incubated at 37°C with shaking (200 rpm —Thermo Scientific MaxQ 6800) until a final OD_600_ of 0·8 was reached. The bacterial cells were harvested by centrifugation and the DNA extraction was performed by the using a Genomic-tip 100/G column (Qiagen) according to the manufacturer’s protocol. The library was prepared according to the instructions of the “Native barcoding genomic DNA (with EXP-NBD114 and SQK-LSK109)” procedure provided by Oxford Nanopore Technology. Sequencing was performed on a MinION Mk1C apparatus (Oxford Nanopore Technologies). The genomic sequences of the isolates were assembled from long and short reads, with a hybrid approach and Unicycler^7^ v0.4.8. A polishing step was performed with Pilon^12^ v1.23 to generate a high-quality sequence composed of chromosomal and plasmid sequences. The sequences are available via the following GenBank accession numbers OR797034 (pPan376-IncN), OR797033 (pPan86), OR797031 (pPan61-IncI1), and OR797032 (pPan61-IncN).

Snippy v4.6.0 (<https://github.com/tseemann/snippy>) was used for mapping all C2 isolates' genomes to pPan376-IncN, pPan86, pPan61-IncI1, and pPan61-IncN using minimum coverage of 4 and base quality of 25. Snippy-core v4.6.0 was used to generate a multiple sequence alignment, and snippy-clean v4.6.0 to replace non-ACTG characters with “N”. Coverage was calculated using SAMtools^13^ v1.14, and presence of plasmid was considered with over 50% coverage. A maximum likelihood tree was constructed for pPan61-IncN (alignment length = 47,695) and pPan376-IncN (alignment length = 44,651) with isolates that had >50% coverage using RAxML-NG^14^ v.1.0.3 with 100 bootstrap replicates to assess support.

### Temporal reconstruction

To estimate the linear relationship between root-to-tip divergence of the input phylogeny and the isolation date the maximum-likelihood tree (Figure 2) of 837 isolates and the corresponding isolate collection year were imported into TempEst^15^. TempEst was run using the “heuristic residual mean squared” and “best fitting root” parameters. Root-to-tip distances were plotted over time, for each *S*. Panama clade and a linear regression analysis was carried out in R. The nucleotide substitution analysis of the four *S*. Panama clades can be found in Table S2.
